# All-atom Molecular Dynamics model for mycobacterial plasma membrane

**DOI:** 10.1101/788299

**Authors:** João L. R. Scaini, Adriano V. Werhli, Vânia R. de Lima, Pedro E. A. da Silva, José Rafael Bordin, Karina S. Machado

## Abstract

Phosphatidyl-myo-inositol mannosides (PIMs) are an essential component of the cell envelope and the most predominant at the inner membrane (IM) of *M. tuberculosis*. In this work, we propose an Molecular Dynamics (MD) *M. tuberculosis* IM model composed of PIM_2_ lipids. The study was divided in three parts: influence of the temperature in the PIM_2_ membrane stability, self-assembly abilities of the PIM_2_ lipid and the behavior when a trans membrane protein is inserted in PIM_2_ membrane. Our results show that the model is able to reproduce the gel phase observed at 310 K and the transition to a fluid phase at 328.15 K. Also, the spontaneous self-assembly of randomly distributed lipids in a vesicular aggregate was observed. Finally, we observe that the PIM_2_ membrane is more stable than DPPC membranes when a Tap protein is inserted. Once Tap eflux pump is related to multidrug resistance of *M. tuberculosis*, this result indicated that the use of the proper lipid model is essential to the proper depiction and modeling of these systems.

**Graphical TOC Entry:** 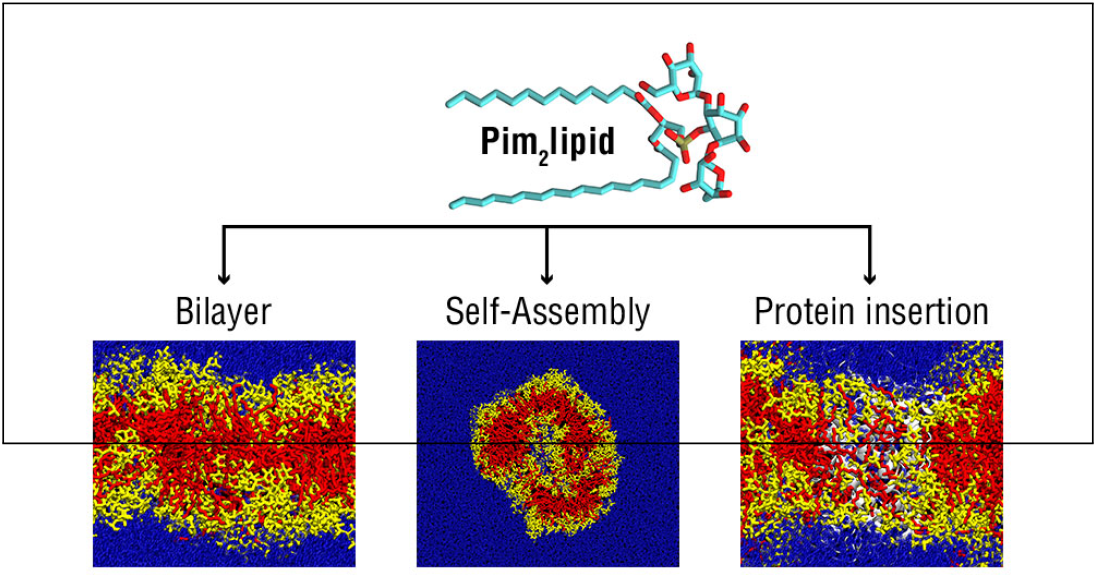

## Introduction

Tuberculosis (TB) is an important infecto-contagious disease caused mainly by *Mycobacterium tuberculosis* being a major cause of mortality and morbidity, specially in developing countries. It was estimated that TB have caused 1.6 million deaths in 2017. The incidence per year for TB has growing since 2013. In 2017, 10 million new cases were estimated. One of the factors attributed to this increase is the emergence of multidrug-resistant (MDR) TB, which was estimated to correspond to 558.000 new cases in 2017. Furthermore, the proportion of MDR-TB cases with extensively drug resistant (XDR) TB has increased, reaching 8.5% in 2017.^1^ In addition to the acquired resistance, *M. tuberculosis* presents high degree of intrinsic resistance related to its cellular ultrastructure. One of the major reasons for the intrinsic resistance to therapeutic agents, and also host defense mechanisms, is the unusual lipid-rich robust and impermeable cell envelope.^2^

*M. tuberculosis* is covered by a complex envelope, composed by the inner plasma membrane (IM), the peptidoglycan-arabinogalactan complex, and the outer membrane (OM) that is covalently linked to the arabinogalactan.^3^ Lipids that are unique to the species or to mycobacteria compose this envelope. The abundance and biological importance of these lipids reflect in extensive studies to elucidate their structures and functions.^4^

Phosphatidyl-myo-inositol mannosides (PIMs) are unique glycolipids abundant in both IM and OM of all mycobacteria, being structural components of the cell envelope. Structurally, PIMs are consisted of a phosphatidyl-myo-inositol (PI) group containing mannosides esterified to the inositol ring.^5^ The PIM family comprises phosphatidyl-myo-inositol mono-, di-, tri-, tetra-, penta-, and hexamannosides, with up to four acyl chains. The two most abundant classes in *M. tuberculosis* are Phosphatidyl-myo-inositol dimannosides (PIM_2_) and Phosphatidyl-myo-inositol hexamannosides (PIM_6_).^6^

Diacyl phosphatidyldimannoside (Ac_2_PIM_2_) is the most abundant lipid in the mycobacterial IM, accounting for up to 42% of all lipids extracted. Other PIMs such as AcPIM_2_, AcPIM_4_, Ac_2_PIM_4_, AcPIM_6_ and Ac_2_PIM_6_ account to up to 26%.^3^ Furthermore, they are precursors of the lipoglycans (lipomannan and lipoarabinomannan), which are multiglycosylated forms of PIM_2_.^7,8^ PIMs and lipoglycans are important virulence factors in *M. tuberculosis* and may modulate to host-pathogen interactions. ^5^

The physicochemical characterization of biological membranes in fundamental to understand their physiological role. ^9^ Elucidation of the mechanisms responsible for the distinct structural conformations and morphologies, and the fusion and dissolution is essential to understand a wide range of biological processes, as for example virulence. ^10,11^ To this end one of the most employed approach to understand the behavior of cell membranes is Molecular Dynamics (MD) simulations. This *in silico* method allow a complete control on the system properties and has been extensively applied to provide information on both their spatial organization and temporal dynamics.^12–14^

Recent MD studies about mycobacterial membrane includes membranes based in DPPC, DOPC, POPG, POPE.^15–17^ Nevertheless, there are no specific membrane molecular model with such uncommon and unique lipids as the observed in the *M. tuberculosis*. Therefore, it would be important to develop a mycobacterial membrane model to allow structural studies that took in consideration its actual lipid composition. As cited before, PIM_2_ is one of the most abundant PIMs, and is precursor of lipoglycans and the other PIMs with higher amount of mannosides. In this way, the main objective of this study is to propose a *M. tuberculosis* IM model made of a PIM_2_ bilayer for MD simulations, analyze its behavior and viability.

In this sense, this study was performed in three parts. First we investigate the temperature effect on a PIM_2_ bilayer to validate it in comparison to its natural behavior. We considered 310 K (36.85 °C) as the normal lungs temperature – where a gel phase is expected – and 328.15 K (55 °C) – the phase transition temperature for mycobacterial membrane. ^18^

The second part of this study aimed to analyze the occurrence of spontaneous aggregation of the lipid molecules forming a bilayer structure^19^ of PIM_2_, as another form of validation of the model. Finally, we studied the behavior of the PIM_2_ bilayer with insertion of a protein that is naturally inserted on *M. tuberculosis* IM and is associated to MDR-TB, the Tap efflux pump.^20,21^

The remaining of the paper is organized as follows: in section 2 we discuss the model and details of the simulation method; in section 3 the results and the main discussions; in section 4, the conclusions are listed.

## The PIM_2_ Model and Simulation Details

### PIM_2_ Bilayer at different temperatures

To obtain the two-dimensional structure of the PIM_2_ molecule we use the Lipid Maps^22^ (LM ID: LMGP15010062). ^4^ The three-dimensional structure was then obtained using Avogadro 1.2.0^23^ minimization. The topology was obtained from the Automated Topology Builder (ATB)^24^ from the submission of the three-dimensional structure, and was manually edited in order to ensure that the names of atoms corresponds to the parameters of the "Berger lipids"^25^ modification for GROMOS96 53a6 force field.^26^ The electrostatic interaction was handled using the Particle Mesh Ewald method.

Molecular Dynamics simulations were performed using the GROMACS 2016.4 package.^27,28^ GROMACS editconf was used to create a 128 lipid bilayer perpendicular to the *z*-axis. A short 50 step minimization and a short 250 step production phase were performed to make the structure flexible. Then, two boxes of water model SPC with a thickness of 1.0 nm were added to both sided of the bilayer solvated the system. Once the system was already neutral, no ions were added.

The solvated system went through energy minimization (EM), equilibration and production. The equilibration was performed with restraints on lipid phosphorus in the *z*-axis, preventing them from leave the position vertically. The equilibration consisted of two phases. The first was a 10.0 ns NVT simulation for temperature adjustment (EQ 1), where the temperature was fixed in 310.0 K or 328.15 K using the velocity rescale. The second was a 10.0 ns NPT simulation (EQ 2) keeping the temperature fixed in 310.0 K or 328.15 K by the Nosè-Hoover thermostat and the pressure fixed in 1.0 bar by the Parrinello-Rahman semi-isotropic barostat – the pressure in the *z*-direction was not fixed. Finally, the system went to a 100.0 ns NPT simulation for the production phase. Here no restraints were applied, and the same temperature and pressure control from the equilibration phase were used.

### PIM_2_ Bilayer formation

MD simulations were performed to test if PIM_2_ molecules would spontaneously aggregate in a bilayer structure. The initial system was a box with 12.0 nm *×* 12.0 nm *×* 6 nm box in the *x*, *y* and *z* directions with 128 randomly distributed PIM_2_ molecules. The box was then fully solvated. The simulations was conducted with the same conditions as the bilayer MDs, and the same steps: energy minimization, 10 ns equilibration for temperature coupling (EQ 1), 10 ns equilibration for pressure coupling (EQ 1 and EQ 2), and 100 ns production. The temperature was fixed in 310.0 K.

### PIM_2_ bilayer with Tap protein inserted

To test the bilayer as an IM model, we performed a MD simulation with a Tap protein^29,30^ inside the PIM_2_ bilayer. The Tap model used was proposed in a previous study of our group.^17^ The same software and force field of the previous sections were employed. The PIM_2_ bilayer final structure from the 100.0 ns 310.0 K MD simulation was used for the insertion. To compare the stability of Tap protein inside of the PIM_2_ bilayer, a simulation with Tap inserted in a dipalmitoylphosphatidylcholine (DPPC) bilayer was performed, following the same protocol. We used coordinates and topology of a previously published stable and flexible DPPC membrane.^31^

The insertion of the protein in the lipid bilayer followed the InflateGRO technique^32,33^ modified according to the protocol of Lemkul.^34^ Briefly stated, it consists in inflating the bilayer area in the *xy*-plane by 4 times, and then deflate the area for 0.95 times, with an EM after each deflation. This allows a a more natural accommodation of the lipid bilayer around the protein. The water from the lipid bilayer is removed in the InflateGRO procedure. In this way, the system was then solvated in the same way as described above. Here we added 5 chlorine ions in the water to neutralize the charge of the system. Next, the system went through EM and equilibration phases with the same configurations as previous section, adding now restraints for the heavy atoms of the protein. This allows shaping the lipid bilayer around the protein. Finally, the same 100 ns production phase at 310 K and 1 bar was performed to analyze the protein inside the distinct bilayer membranes.

## Analysis

GROMACS provided the calculation for some analysis that was plotted using R 3.1.3 and R studio 0.99.491 softwares.^35,36^ The Root-mean-square deviation (RMSD) was calculated at each 2 ps along the entire MD simulations to determine the mobility of the bilayer in different temperatures and with Tap protein. In addition, the RMSD of Tap protein inserted in the bilayer was calculated. The density profile of different atom groups (water, lipid head groups, acyl chains, and protein, when present) was calculated along the *z*-axis to have an overview of the distribution. The deuterium order parameters of the acyl chains of the bilayer were calculated in order to determine how rigid the bilayer is. The lateral diffusion of the lipid bilayer was calculated to determine its fluidity. The thickness of the membrane was calculated using GridMAT-MD^37^ and plottet usind Matplotlib.

## Results and Discussion

The PIM_2_ structure and its three-dimensional structure are shown in the Figures 1a and 1b. The results were described and discussed as follows: First the influence of the temperature in the PIM_2_ bilayer by the comparison of the behavior at 310.0 K and 328.15 K. Then, the bilayer formation MD was described. Finally, the PIM_2_ bilayer was studied with Tap insertion, comparing the MD with Tap insertion with the MD of the bilayer without protein. It is important to note that, for several lipids, computational molecular dynamics data are closest to experimental data when they are in the gel phase.^38^ Therefore, the bilayer formation process and the effects of Tap were studied at 310 K.

**Figure 1:**
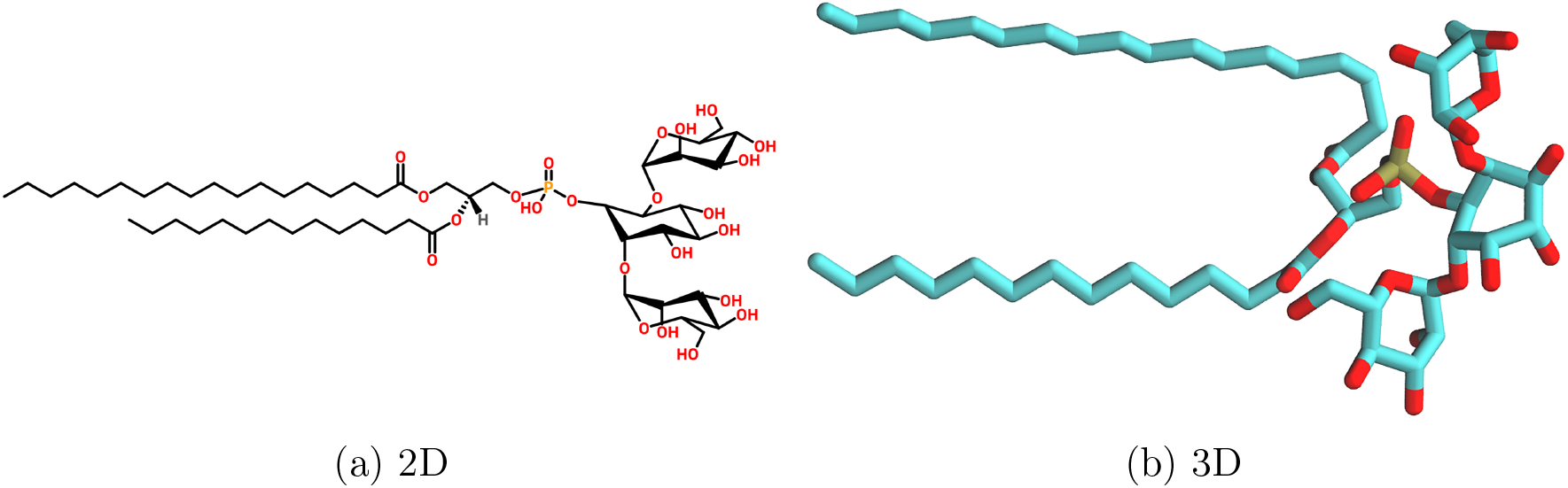
Structure of PIM_2_.

### PIM_2_ bilayer at different temperatures

To validate our model we compare the structural differences of the PIM_2_ bilayer through MD simulations at the lungs temperature (310 K) and at the phase transition temperature (328.15 K). To this end five different properties were analyzed: partial density, RMSD, deuterium order parameters, thickness and lateral diffusion.

### Partial density

The partial density was calculated to analyze the distribution of three atom groups (lipid head groups, lipid acyl chains and water) in the PIM_2_ bilayer system and the temperature effect on it. Both PIM_2_ at 310 K (Figures 3a and 2a) and at 328.15 K (Figures 3b and 2b) shared similar distributions of density. Water remain in the extremes of the *z*-axis and did not penetrate toward the hydrophobic acyl chains – water molecules interacts with the hydrophilic lipid head groups. The acyl chains are denser at the center in both simulations, however at 328.15 K (Figure 3b) they occupy a larger region, with one side denser that the other. At the phase transition temperature, an increase of molecular lateral and rotational diffusion is expected, as well as the intermolecular motion around C-C bonds. Thus, the acyl chains trans conformations will be changed to a gauche one (120 rotation of the simple C-C bond).^39,40^ The increase of molecules at gauche conformation may justify the behavior observed in the model at 328.15 K. Once the PIM_2_ molecule model employed in this study (Figure 1) has different acyl chains (C 18:0 and 14:0), the presence of gauche conformation can highlight the density difference and asymmetry. Thus, our results agrees with the expected behavior: the PIM_2_ bilayer has a more disordered structure at the phase transition temperature and, consequently, is thicker, as will be further discussed in details in the thickness analysis section.

**Figure 2:**
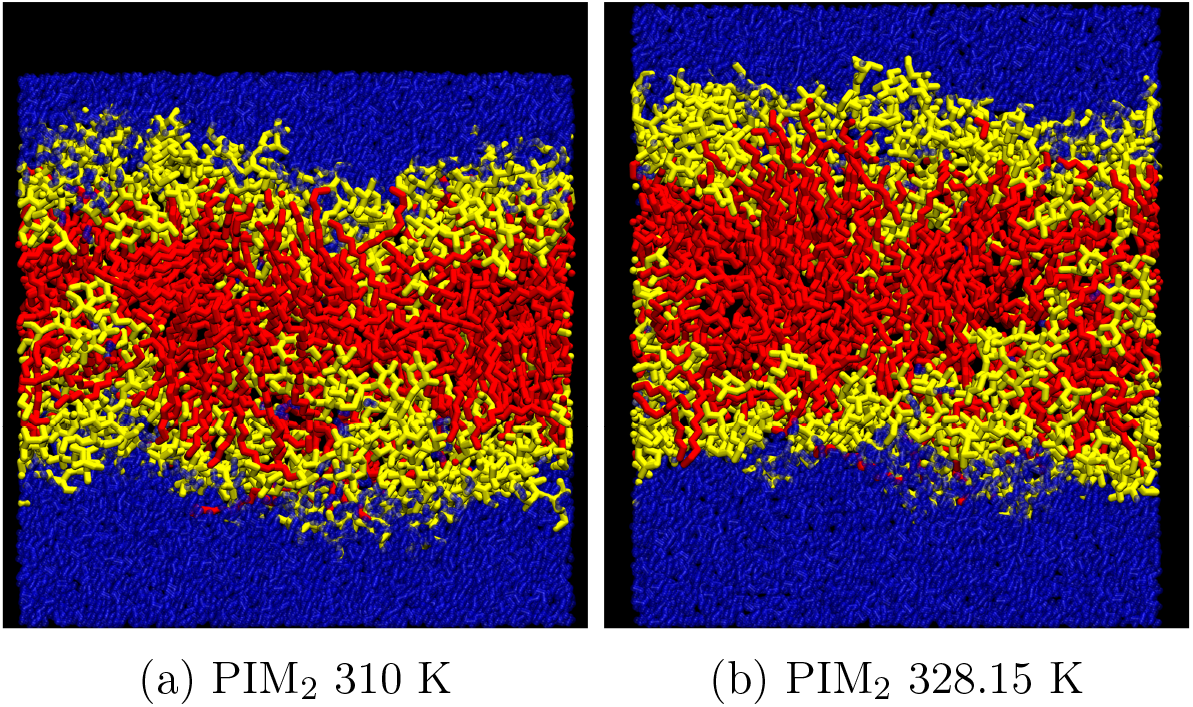
Final structure of PIM_2_ bilayer in three different MDs. (a) bilayer at 310 K, (b) bilayer at 328.15 K

**Figure 3:**
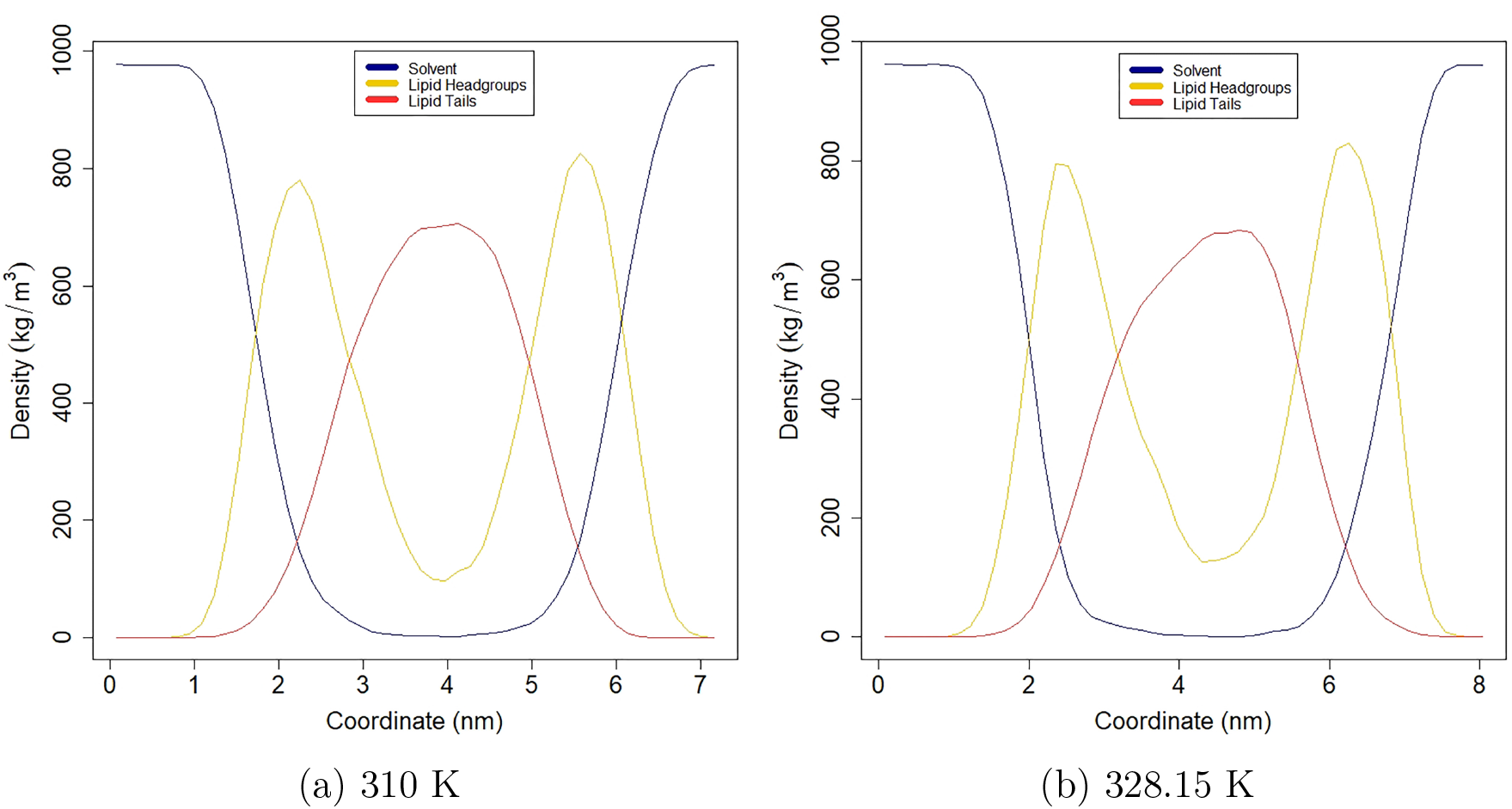
Partial density of PIM_2_ bilayer at different temperatures. *y*-axis represents the density and the *x*-axis represents the coordinates.

### RMSD

The RMSD of the PIM_2_ bilayer was calculated to analyze its stability and the influence of different temperatures on it – it is shown in the Figure 4. PIM_2_ bilayers at 310 K has a lower RMSD that stabilized at 50 ns, staying between 0.95 and 1 nm, which shows the stability of the structure. The lipid bilayer stability is expected as the *M. tuberculosis* membrane exists at this temperature in the lungs and therefore it should be ordered. PIM_2_ at 328.15 K showed a higher RMSD, reaching 1.2 nm at the end of the simulation, and did not stabilize. Kirsch and Böckmann^41^ showed that a phospholipid bilayer have a reduced stability close to the phase transition temperature. Therefore, this higher and more unstable RMSD from PIM_2_ at 328.15K agrees what is expected for the liquid state, indicating a disordering process at phase transition temperature.

**Figure 4:**
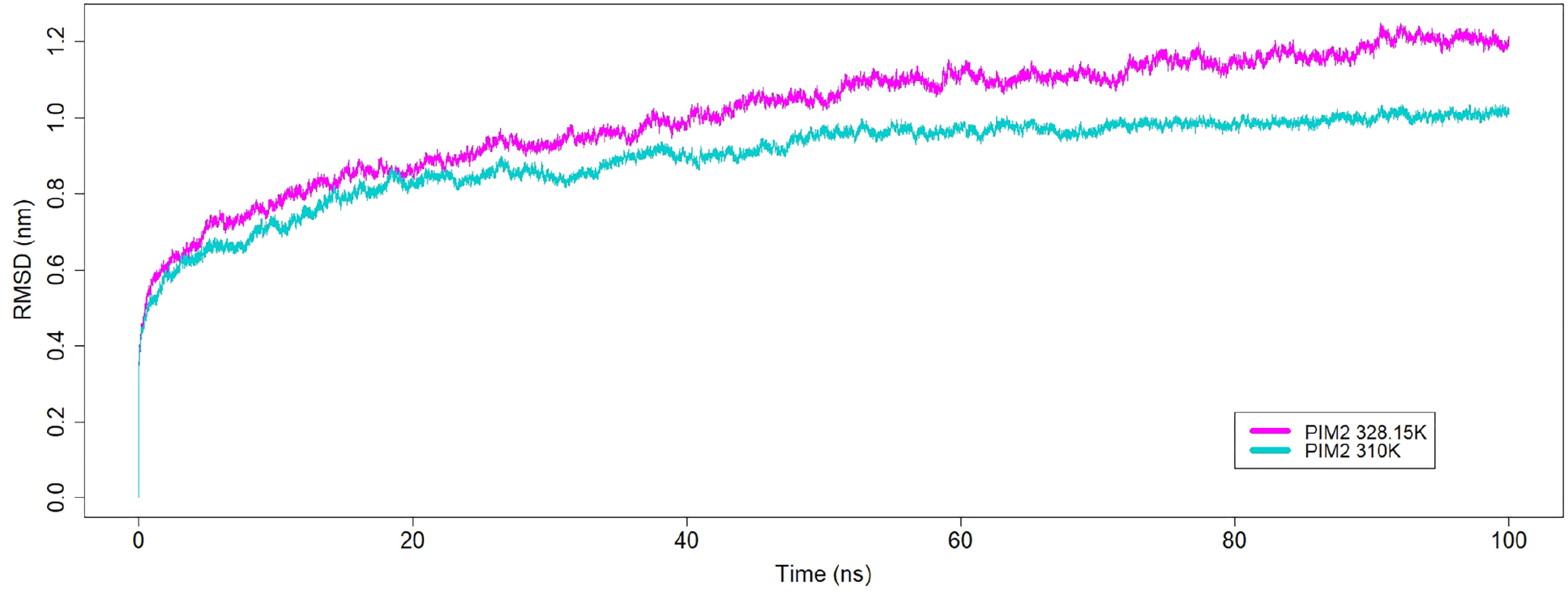
RMSD of PIM_2_ bilayer in the 100.0 ns MD simulations at different temperatures. *y*-axis represents the RMSD in nm and the *x*-axis represents the time in ns.

### Deuterium order parameters

Deuterium order parameters were calculated to analyze how ordered were the acyl chains of the PIM_2_ bilayer system and the temperature influence on it, shown in Figure 5. Both chains, in both temperatures, are more ordered at the beginning of the chain, closest to the head groups. It is expected, once the acyl chains are more prone to move. Both acyl chain 1, shown in the Figure 5a, and acyl chain 2, Figure 5b, are less ordered at 328.15 K. Once more ordered acyl chains are indicative of a gel state, it is a coherent result that indicates a fusion at 328.15K. For lipid acyl chains, the correlation of time and order parameters related to the molecular overall motion are the same for all methylene and methyl groups. However, chain conformation and trans gauche isomerism rate show a flexibility gradient, raising from the first chain methylene to the terminal methyl group. As trans-gauche isomerism increases close to the melting temperature, it is expected a reduction in the S_cd_ values related to the model at 328.15 K, compared to the values observed at 310 K.^42^

**Figure 5:**
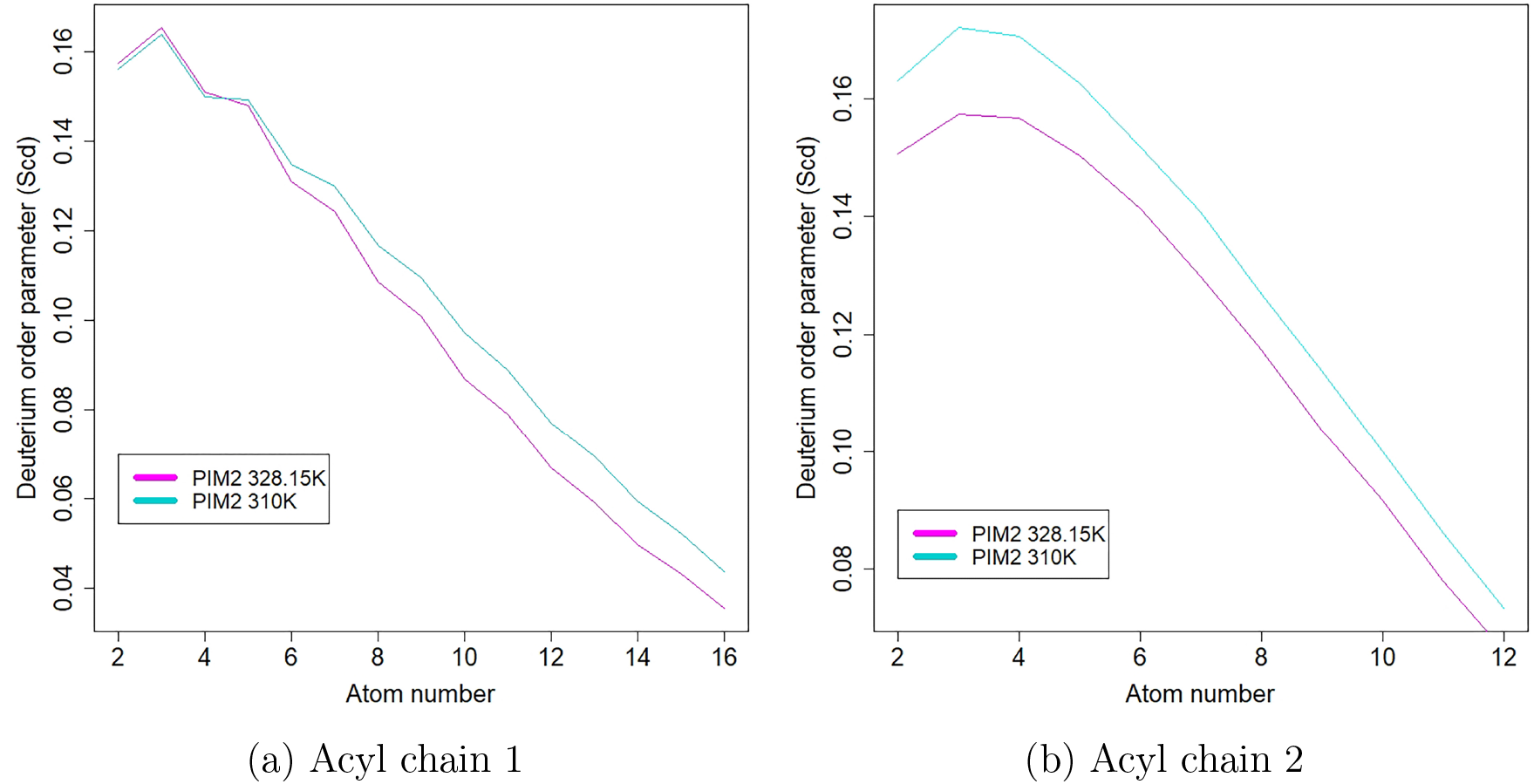
Deuterium order parameters of PIM_2_ bilayer acyl chains at different temperatures. *y*-axis represents the deuterium order parameter and the *x*-axis represents the atom number.

### Thickness

The thickness of PIM_2_ bilayer system, shown in Figure 6, was evaluated to analyze the difference of its thickness at different temperatures. This graphic shows the thickness according to region. The PIM_2_ bilayer at 310 K showed 21.5% of its structure with thickness over 4 nm, where the thickest region is 4.42 nm. While the structure at 328.15 K was 63% thicker than 4 nm, with the thickest region of 5.048 nm. Atomistic simulations revealed a minimal thickness at lipid interface close to melting temperature. ^41^ Thus, we can observe that at 328.15K there are more thicker regions, which is expected, since the phase transition temperature leaves the membrane disordered. Here it is important to note that membrane thickness may also correlate with permeability, which is also a natural characteristic at phase transition. ^43^

**Figure 6:**
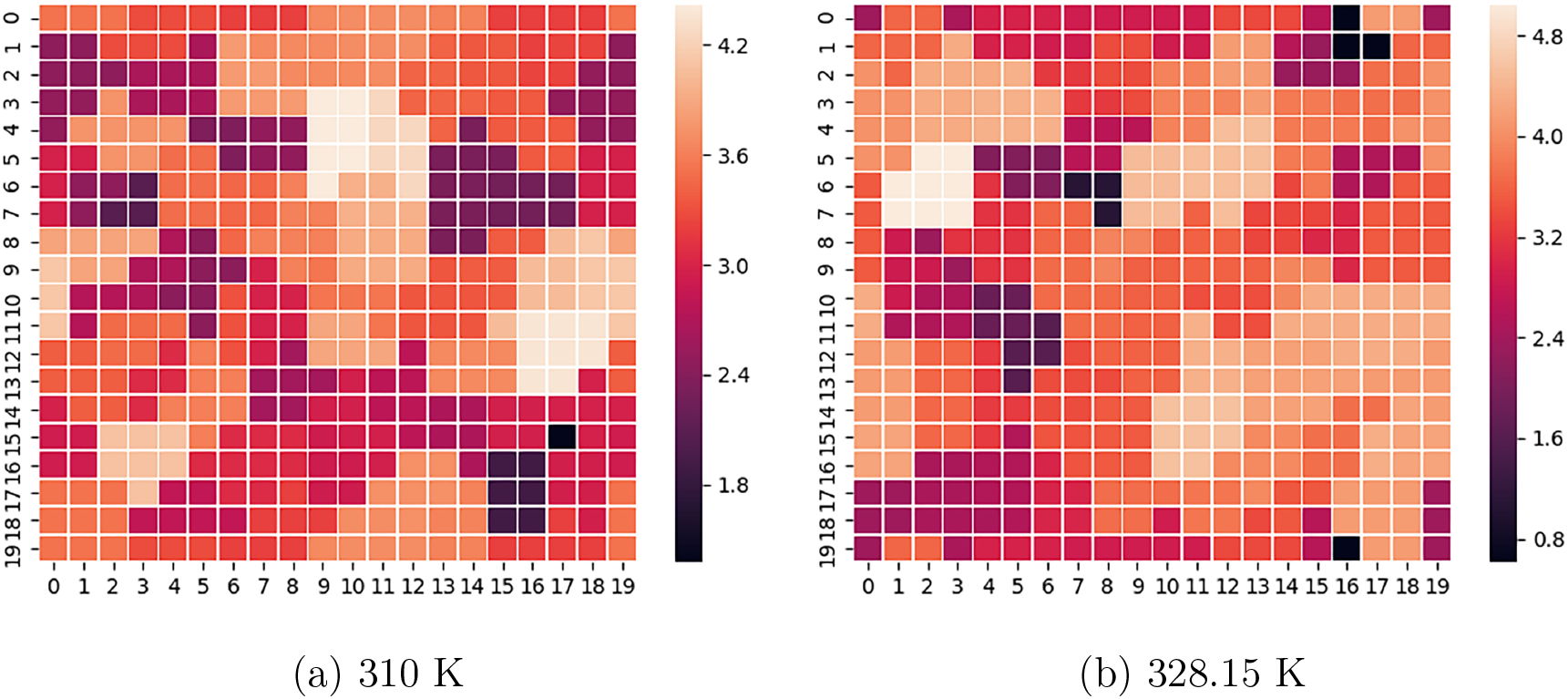
Bilayer Thickness of PIM_2_ bilayer at different temperatures. *y*-axis and *x*-axis are a 20 x 20 points grid box representing the PIM_2_ bilayer area. The *z*-axis is represented by a color gradient indicating the thickness (nm), where darker colors are thicker regions.

### Lateral Diffusion

The MSD of the lateral diffusion coefficient of the P atom was calculated to analyze the lateral diffusion in PIM_2_ and the temperature influence on it (Figure 7). The lateral diffusion coefficient of PIM_2_ at 328.15 K reaches around 1.5*×*10^*−*5^ cm^2^/s, while at 310 K it reaches 0.7*×*10^*−*5^ cm^2^/s. For entire dimyristoyl phosphatidylcholine molecules the lateral diffusion value is 17*×*10^*−*8^ cm^2^ in the fluid phase. When the lipid is in the gel phase, this value is equivalent to 7*×*10^*−*8^ cm^2^/s (considering DMPC melting temperature equivalent to 297.15 K). The typical increase of lateral diffusion as the membrane transits from gel to fluid phase was also observed to PIM_2_ bilayer. It is coherent to what is expected, once the phase transition temperature makes the membrane more flexible, therefore increasing the lateral diffusion coefficient.

**Figure 7:**
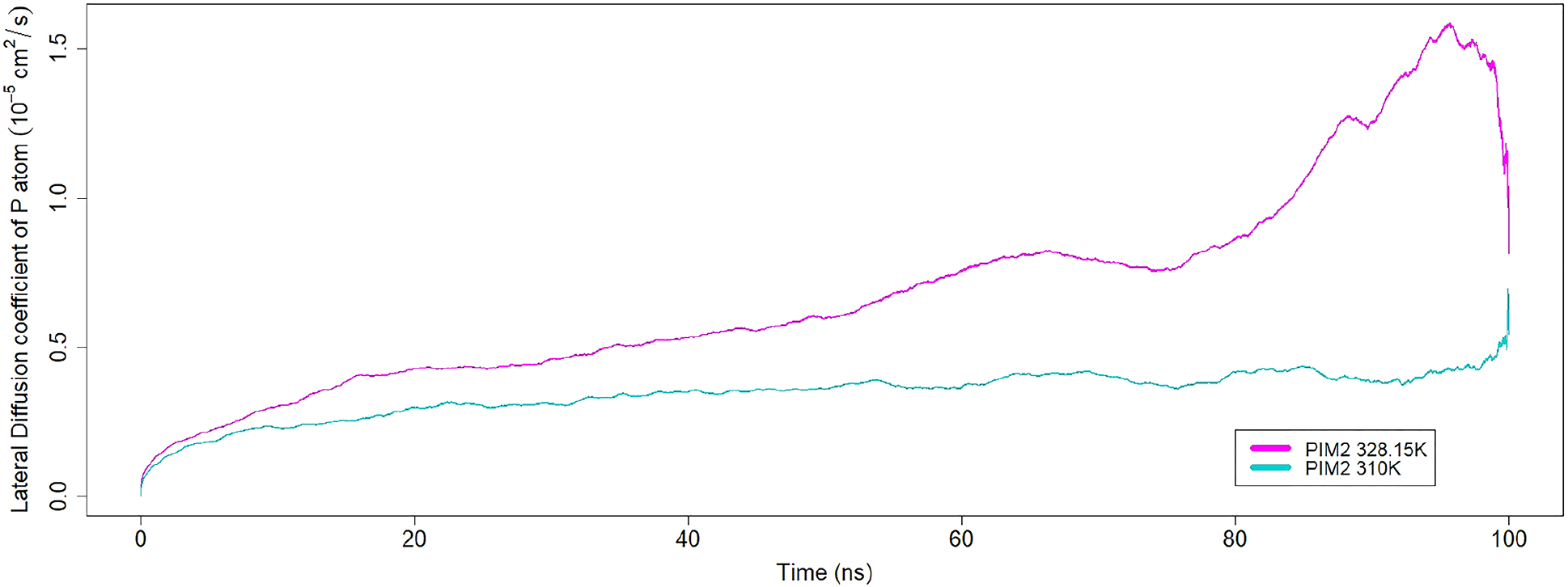
Lateral diffusion coefficient of the PIM_2_ bilayer in 100 ns MD simulations at different temperatures. *y*-axis represents the RMSD in nm and the *x*-axis represents the time in ns.

### PIM_2_ Bilayer Formation

In order to test the lipid property of aggregation in water of the PIM_2_ molecule we performed a simulation with the lipids starting at a random positions. Snapshots of the aggregation process can be seen at Figure 8. It started with 128 lipids randomly distributed in the box (Figure 8a). During the first 10 ns equilibration (EQ 1), two micellar and one vesicular aggregates were observed at 5 ns (Figure 8b), and at the final structure of this phase two large aggregates were observed: a vesicular one and a spherical micelle, as shown in Figure 8c. At the end of the second 10 ns equilibration a single vesicular aggregate is observed, as shown in Figure 8d. Finally, at the end of the 100 ns production phase, there is the final vesicular aggregate, with only one pore-like defect. Studies of charged species permeation in membrane show the formation of a water process finger defects in the membrane.^43^ Other explanation for the observed defect is the transient pore formation mechanism, which stability depends to the proton transport through membrane. ^44^ Also, systems with self-assembly properties may require even longer simulation to relax to the final configuration.^45^

**Figure 8:**
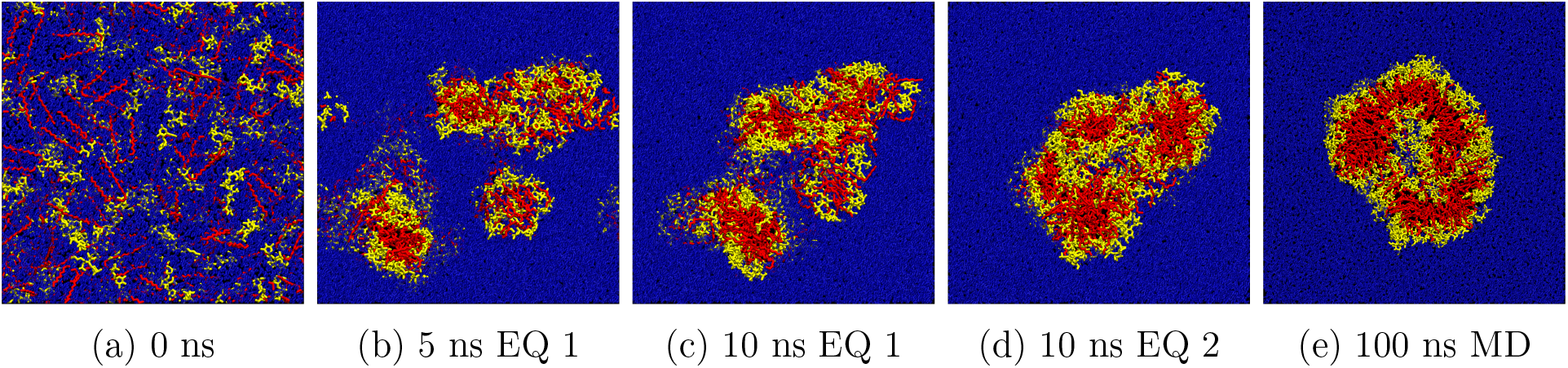
Snapshots of Molecular Dynamics simulation of PIM_2_ aggregation in water. Lipid head groups (yellow); acyl chains (red); water (blue). (a) is the first frame. (b) is a snapshot from 5 ns of the 10 ns Equilibration step (EQ 1). (c) is the final structure of the first equilibration. (d) is the final structure of the second equilibration (EQ 1. (e) is the final structure of the production MD.

### PIM_2_ bilayer with insertion of Tap

In order to validate the model, we analyzed its behavior with a transmembrane protein found in *M. tuberculosis* IM, Tap efflux pump. For this end, we compare the structural differences of PIM_2_ bilayer during the MD simulation with Tap protein inserted (Figure 10) with the MD simulation of the PIM_2_ bilayer at 310 K previously discussed (Figure 2a), once both simulation were performed at the same temperature.

**Figure 9:**
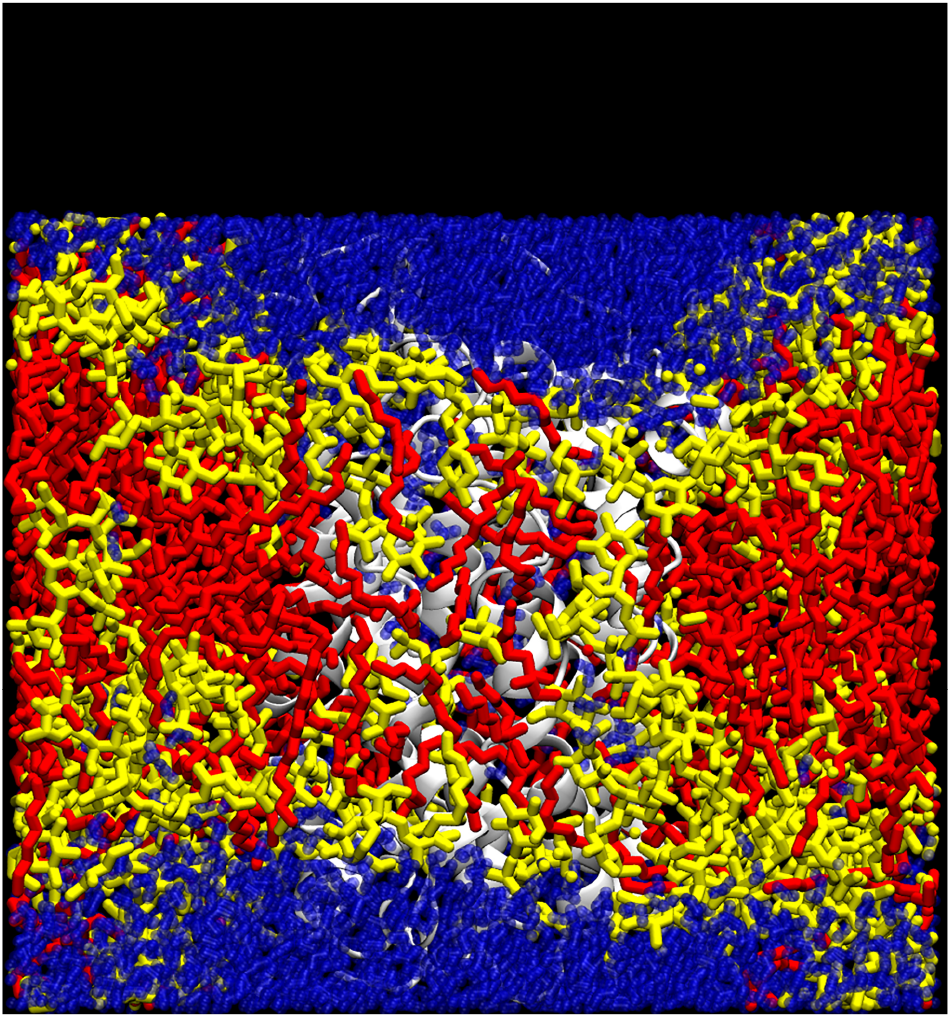
PIM_2_ with Tap

Figure 10: Final structure of PIM_2_ bilayer with Tap protein insertion. Lipid head groups (yellow), Lipid acyl chains (red), water (blue) and protein (white)

### Partial Density

The partial density shown in Figure 11 was calculated to analyze the distribution of four atom groups (lipid head groups, lipid acyl chains, protein and water) in the PIM_2_ bilayer system and the effect of Tap protein insertion on it. The lipids and water atoms had a similar density distribution to the PIM_2_ bilayer simulation without protein. Water also stayed in the extremes of the *z*-axis, did not penetrate toward the acyl chains, and interacted with the head groups. The acyl chains were also dense at the center. However, the head groups and the acyl chains had more intersection in comparison to the PIM_2_ bilayer MD without protein insertion (Figure 3a), where the separation was more clear. That is a consequence from the fact that the lipids were tighter together, showing a less clearly delimited structure. The protein was across the whole bilayer, and is more centered towards coordinate 0 at *z* axis than the other extreme of the *z*-axis.

**Figure 11:**
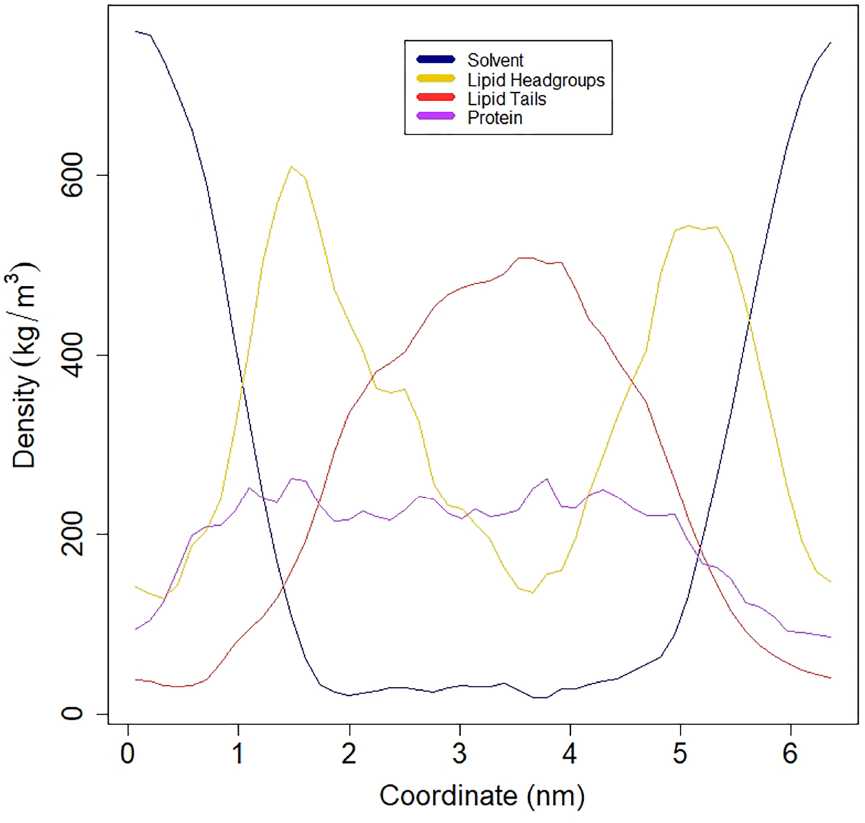
Partial density of PIM_2_ bilayer with Tap protein insertion. *y*-axis represents the density and the *x*-axis represents the coordinates.

### Bilayer RMSD

The RMSD of PIM_2_ bilayer with Tap protein inserted was calculated to analyze the stability of the bilayer and the effect of Tap insertion in it – Figure 12. PIM_2_ bilayer with Tap showed a lower RMSD, but that never stabilized. This lower RMSD is expected and reaffirms the PIM_2_ bilayer as a *M. tuberculosis* IM model, that therefore has transmembrane proteins, like Tap, naturally inserted in it. The instability is a consequence from the fact that the system is more crowded.^46,47^

**Figure 12:**
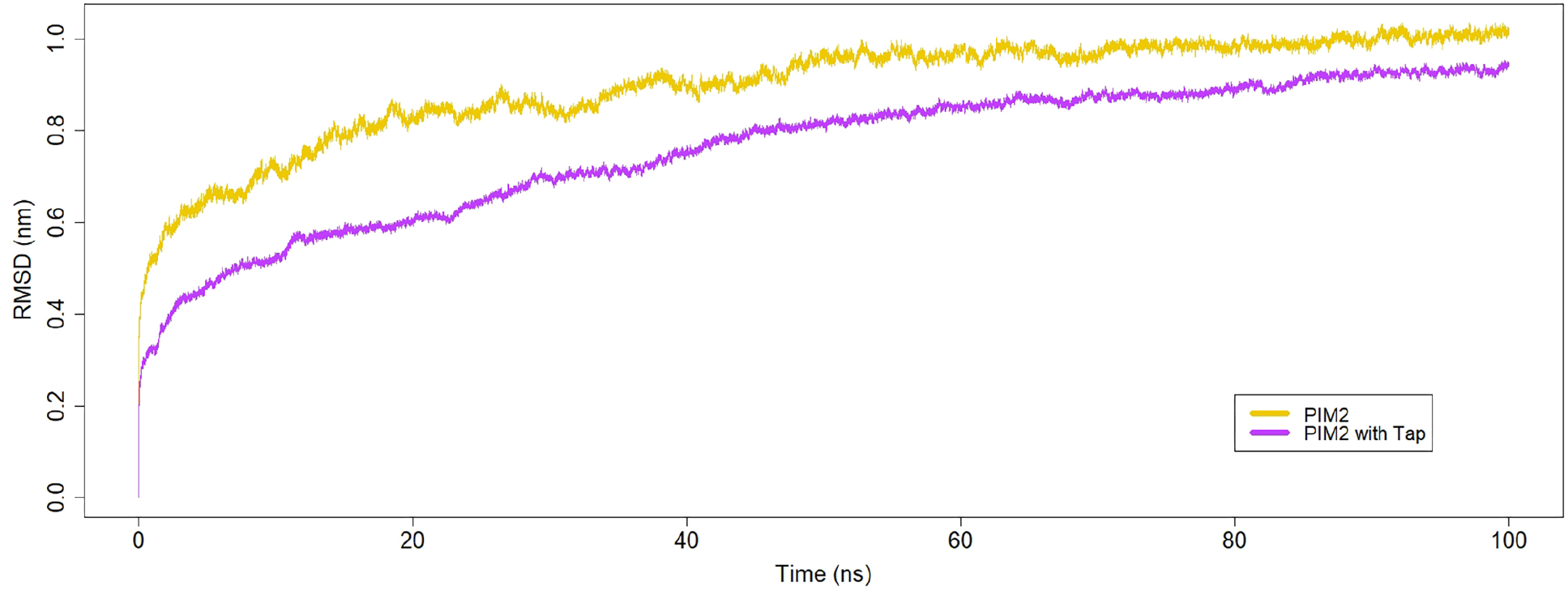
RMSD of PIM_2_ bilayer in 100 ns MD simulations with and without Tap protein insertion. *y*-axis represents the RMSD in nm and the *x*-axis represents the time in ns.

### Deuterium Order Parameters

Deuterium order parameters were calculated to analyze how ordered were the acyl chains of the PIM_2_ bilayer with Tap protein insertion and the influence of Tap insertion on it (Figure 13). A higher deuterium order means a more ordered structure. Both acyl chains have higher deuterium parameters in PIM_2_ with Tap in comparison to PIM_2_ without Tap. This indicated that acyl chains are more organized in the structure with the Tap inserted inside. This indicates a more ordered structure with Tap protein insertion, which reinforces the capability of the PIM_2_ model for *M. tuberculosis* IM.

**Figure 13:**
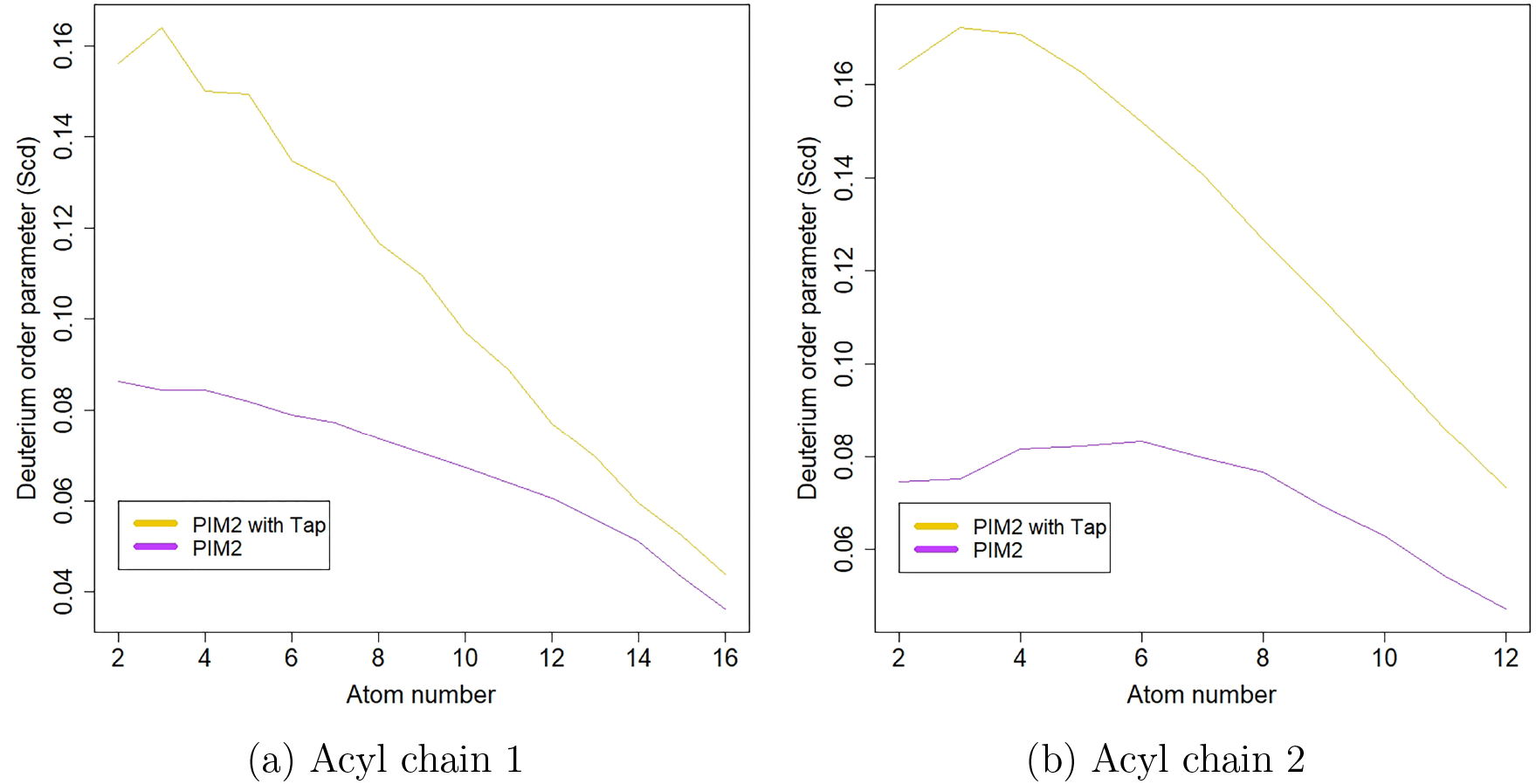
Deuterium order parameters of PIM_2_ bilayer acyl chains with and without Tap protein insertion. *y*-axis represents the deuterium order parameter and the *x*-axis represents the atom number.

### Bilayer Thickness

The thickness of PIM_2_ bilayer system with Tap protein insertion, shown in Figure 14, was calculated to analyze the difference of its thickness and the influence of Tap insertion on it. The PIM_2_ bilayer with Tap had 36.45% of its structure thicker than 4 nm, where the thickest region is 6.154 nm. It is thicker than PIM_2_ bilayer e without Tap insertion, that had 21.5% of its structure thicker than 4 nm and the thickest region is 4.420 nm. These thicker regions can be explained because the system is more crowded, and therefore had to agglutinate more. But it can be observed that the thickest regions of PIM_2_ with Tap are in extreme regions, which are far from the protein. The bilayer structure around the protein is compact, and therefore rigid, near the protein, which is the most important when studying proteins inserted in membranes, and also shows an adequate behavior that reinforces our bilayer as a efficient *M. tuberculosis* IM model.

**Figure 14:**
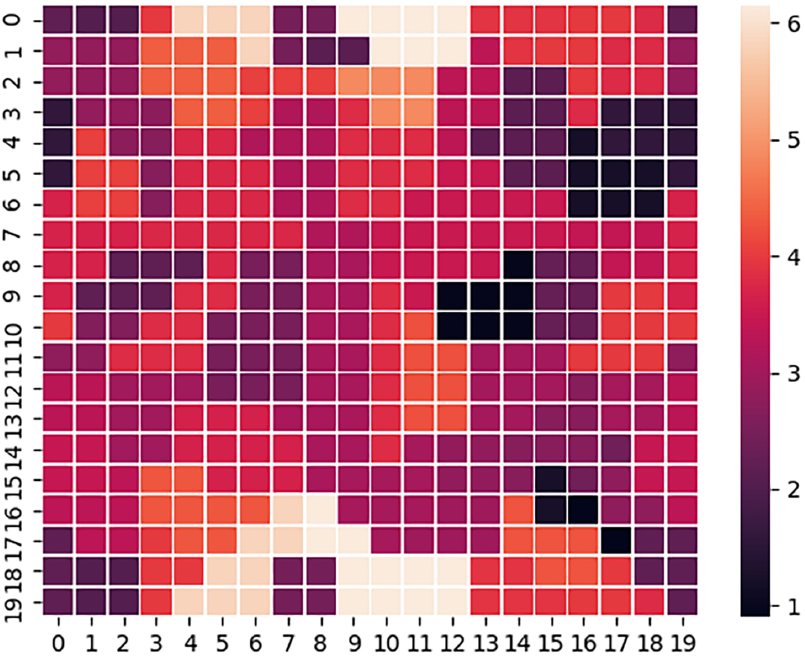
Bilayer Thickness of PIM_2_ bilayer with Tap protein insertion. *y*-axis and *x*-axis are a grid box representing the PIM_2_ bilayer area. The *z*-axis is represented by a color gradient indicating the thickness (nm), where darker colors are thicker regions.

### Lateral Diffusion

The MSD of the lateral diffusion coefficient of the P atoms were calculated to analyze the lateral diffusion in PIM_2_ bilayer system with Tap protein insertion and the influence of Tap insertion on it, as shown in the Figure 15. The lateral diffusion of PIM_2_ with Tap insertion, reaches around 0.4*×*10^*−*5^ cm^2^/s, and is lower than PIM_2_ without the protein, that reaches around 0.7*×*10^*−*5^ cm^2^/s. This indicates that the protein insertion makes the bilayer more rigid. This is the expected, reinforcing our proposition for a lipid model for the *M. tuberculosis* IM.

**Figure 15:**
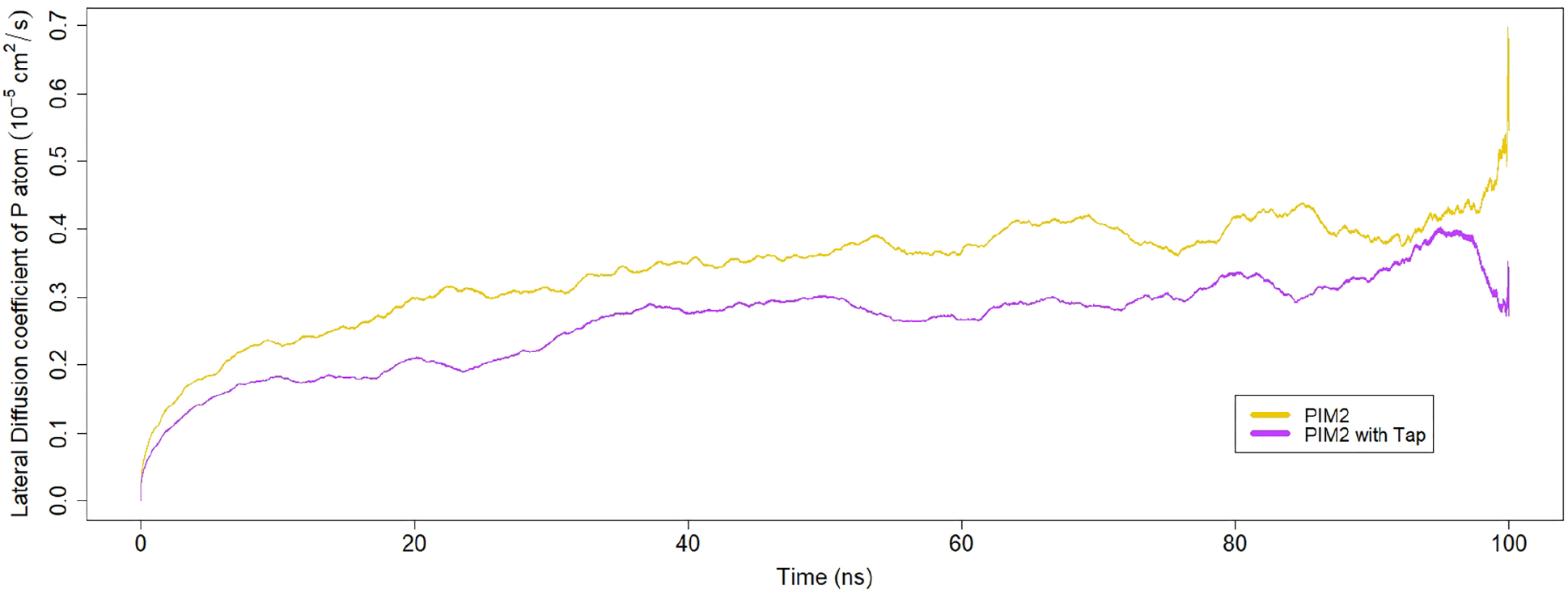
Lateral diffusion coefficient of the PIM_2_ bilayer in 100 ns MD simulations with and without Tap protein insertion. *y*-axis represents the RMSD in nm and the *x*-axis represents the time in ns.

### Protein RMSD

Finally we analyze the stability of Tap inserted in the PIM_2_ bilayer. For this end we use RMSD calculations for the MD simulations of both Tap in PIM_2_ and Tap in DPPC, both at 310 K, shown in the Figure 16. The atoms from the helices were accounted, once they are a more rigid structure than the turns and are the most predominant.^17^ Tap in DPPC had a lower RMSD, but never stabilized, growing to even higher RMSD than Tap in PIM_2_. Tap in PIM_2_ has a higher RMSD at first, but it was stable from the beginning, stabilizing definitely at 40 ns. This stabilization reinforces the PIM_2_ role of anchor lipid, allowing the protein typical from *M. tuberculosis* IM to stabilize, and therefore maintain a functional structure. This result show that the choice of the appropriate lipid to compose the membrane can affect the transmembrane protein behavior. Therefore the lipid type and model should be carefully chosen. This final data reinforces the necessity of models for PIMs lipids.

**Figure 16:**
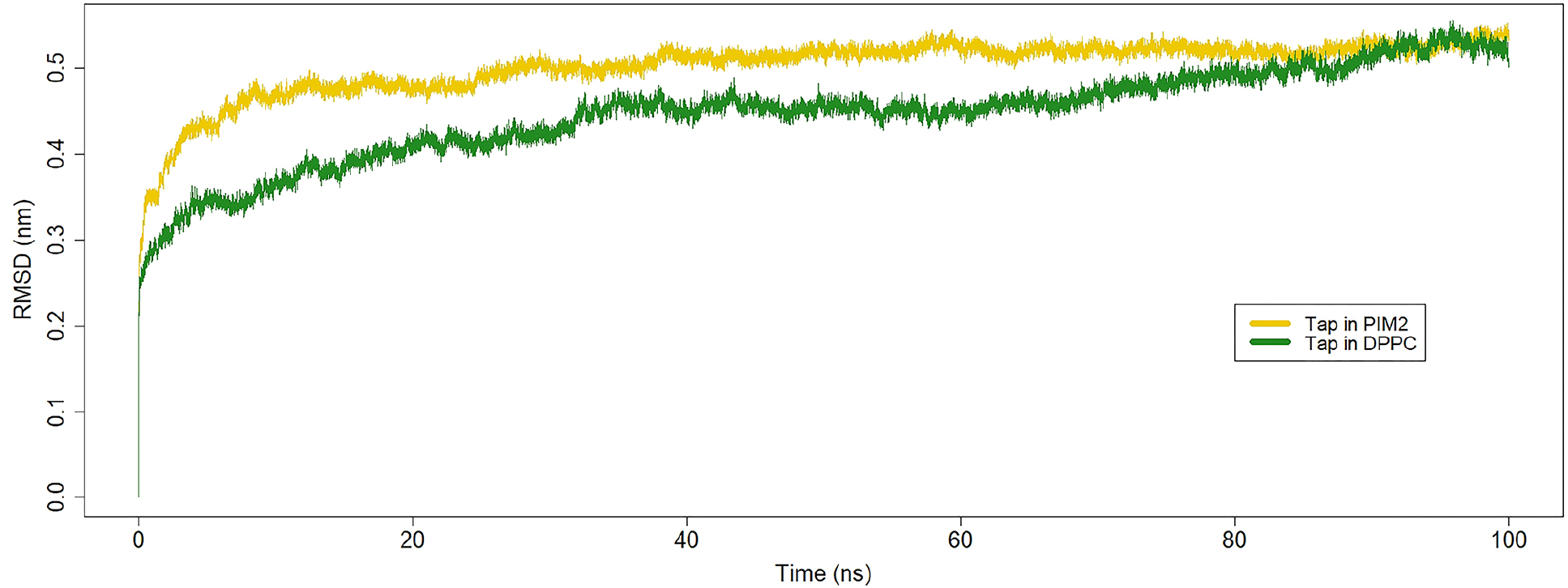
RMSD of Tap Helix during 100 ns MD simulations in bilayers of different lipids. *y*-axis represents the RMSD in nm and the *x*-axis represents the time in ns.

## Conclusion

Is this work we performed the first simulations of PIM_2_ bilayer as a model for *M. tuberculosis* IM. Our results showed that the behavior of our bilayer PIM_2_ model at different temperatures was coherent for lipids in both gel and fluid phases, showing viability to study the temperature dependence of this membrane. This is important since many conditions can be simulated, specially between the natural body temperature and the phase transition temperature of the membrane. Also, a spontaneous aggregation process was observed, which shows the ability of the model to self-assemble in a bilayer.

The behavior of the bilayer with the insertion of Tap protein also highlights the IM model as reliable. The bilayer was even more stable with the insertion of the protein. The protein also reached stability faster in the PIM_2_ bilayer than in a DPPC model. Therefore, we conclude that this model can be used for studies using trans membrane proteins typical of membranes rich in PIM_2_ lipids. Molecular Dynamics studies using Tap has been published, and therefore a model that is adequate for protein insertion is an important step. Also, this model can be employed to study others trans membrane proteins, nanoparticle based drug delivery systems, and others process that occurs in the bacterial membrane.

## Acknowledgement

The authors thank the support from the Brazilian Agencies CNPq, CAPES and FAPERGS. JLRS thanks the Coordenação de Aperfeiçoamento de Pessoal de Nível Superior (CAPES), Finance Code 001. Without the public funding this research would be impossible.

